# Hidden State Genomics: *Graph-Based Analysis of Sparse Auto-Encoder Feature Activity in Genomic Language Models*

**DOI:** 10.64898/2026.05.13.725007

**Authors:** Eliot Kmiec, Samuel O’Brien, Matthew McCoy

## Abstract

Pre-trained genomic language model (gLM) representations have been anticipated to enable enhanced deep learning predictions on several genomics tasks, but current benchmarking has led to questions over what they actually encode. We studied this with mechanistic interpretability on InstaDeep’s Nucleotide Transformer v2 (500M), training sparse autoencoders across all 24 encoder layers to probe latent features. Correlation-based annotation against reference regulatory tracks was inconsistent across layers and insufficient for causal interpretation. We therefore built typed sequence-to-feature knowledge graphs to explore the SAE feature space and compared cisplatin-binding versus non-binding genomic DNA sequence communities by PageRank centrality, validating candidate features with decoder-based interventions and a CNN binding classifier. Interventions showed asymmetric effects: suppressive features could collapse predictive signal, while binding-associated features shifted predictions cumulatively with the presence of other binding-associated signals. Dependency maps further indicated strong local feature sensitivity within sequences. Together, these results provide evidence that gLM representations encode highly granular sequence syntax and conservation patterns, aligning more strongly with tightly coupled molecular interactions and local biophysical constraints than with complex, distributed regulatory logic. Within the scope of our intervention setting, this pattern is consistent with stronger performance on selected molecular tasks and weaker performance on broader regulatory inference, motivating scalable methods for causal feature annotation.

## Background

The success of pre-trained foundation models in natural language processing tasks such as text-generation and language translation has led to the training of a wide array of Genomic Language Models (gLMs) based on Artificial Neural Networks (ANNs) [1, 2]. These models are trained on DNA sequences using a self-supervised training objective to learn generalized representations of the information contained within genomic sequence data. Typically, these training objectives mirror those used in natural language processing, but the specific objective varies by model family: masked language modeling estimates intentionally masked positions from surrounding context, whereas autoregressive or causal language modeling predicts the next token in sequence. Many encoder-based gLMs, including the Nucleotide Transformer models discussed here, use masked language modeling during pre-training. These processes have been shown in other domains to reliably produce information-rich general representations of sequence data that can be utilized for multiple downstream predictive tasks. Thus, gLMs have been expected to yield similar benefits for predictive modelling on tasks where genomic sequence data is utilized [3].

Despite the success of foundation models in other contexts, such as the GPT models that form the basis of ChatGPT, gLMs have struggled to deliver similar performance benefits to some downstream applications in genomics, and often require significant computing resources allocated for fine-tuning in order to achieve desired performance [4, 5]. Due to the complexity of these models, published discourse on the limitations of gLMs currently focuses on performance benchmarking, and mechanisms are largely derived from intuition rather than empirical study. While the ANN architecture that these foundation models have been built on has long been considered a “black box”, significant advances have recently been made toward developing methods to understand their activation patterns and post-training behavior.

In 2024, researchers at Anthropic demonstrated the use of Sparse Auto-Encoders (SAEs) to extract interpretable latent vectors from the activation space of their flagship large language model, Claude [6]. This approach is based on the theory of superposition, which states that the perceptrons in an ANN represent features in the data using combinations of perceptron outputs rather than the activity of a single perceptron. The SAE allows the isolation of features encoded by the model by expanding the latent space and applying a sparsity constraint. SAEs typically achieve this by utilizing a single hidden layer with an L1 sparsity penalty applied, and scaling the hidden layer to a chosen multiple of the original embedding matrix dimensionality. An annotation process is then used to assign meaning to these new feature vectors, also referred to as latents, typically by providing sample activity from the latents to a Large Language Model (LLM) for automated analysis. To validate the labels, steering experiments are performed using the decoder from the SAE to construct embeddings with artificial latents and then fed through the original model to see what effect it has on the output. Two independent groups have so far successfully applied these methods to ESM-2, revealing interpretable features of proteins learned by the foundation model [7, 8].

Applying these methods to gLMs has the potential to address some major questions about their performance. In 2024, Boshar and colleagues compared performance of gLMs and Protein Language Models (pLMs), and found evidence that gLMs such as InstaDeep’s Nucleotide Transformer family of models could be used to predict properties of transcribed proteins with comparable performance to pLMs. Yet, in 2025, Tang and colleagues found that gLMs failed to recognize cell-specific regulatory motifs and some models had worse performance than convolutional neural networks (CNNs) prior to task-specific fine-tuning [9, 4]. This performance gap between the domain in which the models are trained and the domains in which they perform well seems to suggest a misalignment of pre-training incentives and desired feature encoding; however, specific knowledge of what gLMs encode is currently an open research question. Having knowledge of what is encoded by gLMs after pre-training would provide needed insight into the mode of failure that is producing the misalignment, as well as potentially revealing novel biological mechanisms.

### Model Selection and SAE Training

InstaDeep’s Nucleotide Transformer (v2) series of models were selected for this study, as the models are open source, readily accessible via the HuggingFace ecosystem, and have been utilized in multiple previous studies [2]. With the resources we had access to, we trained 72 SAEs on embeddings from all 24 layers of InstaDeep’s NTv2-500m-human-ref encoder (NTv2) using a set of approximately 20,000 sequences from the human reference genome (GRCh38/hg38, GENCODE v49). NTv2 is a traditional encoder-only transformer architecture, and the version we used was pre-trained on 500 million sequences from the human reference genome. All downstream models were trained with the NTv2 weights frozen for consistency. Each SAE is composed of an input layer whose dimensions match the NTv2 embedding dimension, a single hidden layer where the L1 sparsity penalty is applied to an expanded latent space of 8, 16, or 32 times the original embedding dimension; and a single decoding layer which reconstructs the original embeddings. These SAEs were trained using an L1 sparsity penalty of 0.001, and a linear annealing schedule of 100 steps as shown by Simon and Zou to gradually increase the L1 sparsity penalty for training stability [7].

### Limitations of Correlation Scores for Annotation

Use of LLMs to automate the annotation process for mechanistic interpretability studies on LLMs and pLMs is enabled by the natural language domain and readily accessible protein-specific metadata from the Protein Data Bank, respectively [6, 7]. DNA is not a human-readable language, and the functions of individual elements are often dependent on interactions with other elements in the genome, so the annotation process in gLMs is necessarily more complex. Earlier mechanistic interpretability experiments on gLMs have attempted to utilize correlation scores between NCBI reference tracks and SAE latents to label individual features; however, this approach has limitations [1].

Although the correlation approach is an intuitive adaptation, there is no inherent causality in correlations, thus it is extremely difficult to design sound steering experiments to validate annotations generated from correlations. This poses a major epistemic limitation compared to information-theoretic approaches such as attribution maps [10]. Additionally, it is impossible to label anything other than previously understood biology by this method, thereby eliminating one of the anticipated benefits of interpretability study on gLMs.

In our experiments, we also find that correlation between latents and known references varies empirically across layers. We computed Pearson and normalized cross-correlation coefficients for each feature against GRCh38/hg38 RefSeq regulatory element tracks obtained from the UCSC genome browser. Feature signal per token was used to calculate these correlations and identify the most highly correlated feature for each regulatory element track as displayed in *Figure 1*. By plotting the best correlated feature scores for these regulatory elements layer by layer, we demonstrate how likely the latents are to extract the desired information and which layers extract this information better than others. While we do expect natural variations in per layer representations, we generally expect deeper layers of the encoder to capture more information and for the trend to be positively correlated with the layer depth.

**Figure 1:**
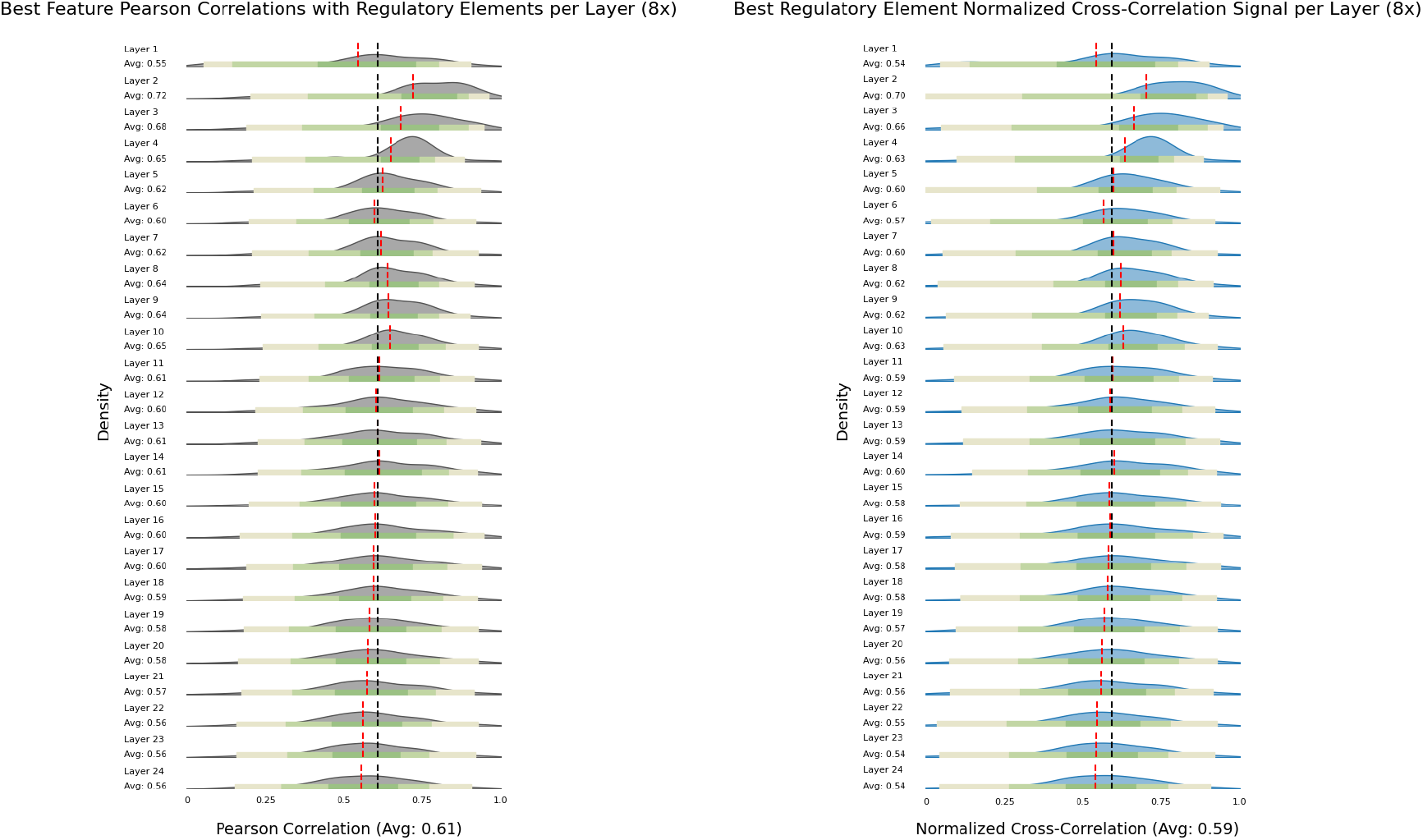
Ridgeline density plot showing distribution of highest correlated features for the NCBI RefSeq regulatory element track set from the UCSC genome browser. Comparison between Pearson metric (left) and Normalized Cross-Correlation (right) to account for phase / amplitude differences. Black dotted lines indicate global average, red dotted lines indicate local layer-wise average. Colored bars represent quartiles.

Contrary to the expected pattern, after the initial spike in the second layer, the SAEs struggle to find latents that capture the labeled regulatory elements. This suggests that either the hypothesis by Tang and colleagues [4] about misaligned pre-training incentives is correct, or that the latents are splitting in ways that favor minute syntax over broad categories of regulatory logic [6]. Some initial evidence for the latter hypothesis can be found in Figure 2, which compares subcategories of regulatory elements belonging to the family of Long Interspersed Nuclear Elements (LINE) and the family of protein binding sequences (PBS). LINEs are a group of highly specific transposons found throughout the human genome, and we see that deeper representations are necessary for the SAEs to extract information related to them, whereas PBSs are a large category of general functionality which retains the negative trend across layers. However, we observed similar trends and average correlation differences of less than 0.02 across different SAE expansion sizes, where we would expect splitting of latents to be more significant. If splitting of latents is responsible for this phenomenon, it cannot be determined from correlation alone, motivating the causal and graph-theoretic approaches that follow.

**Figure 2:**
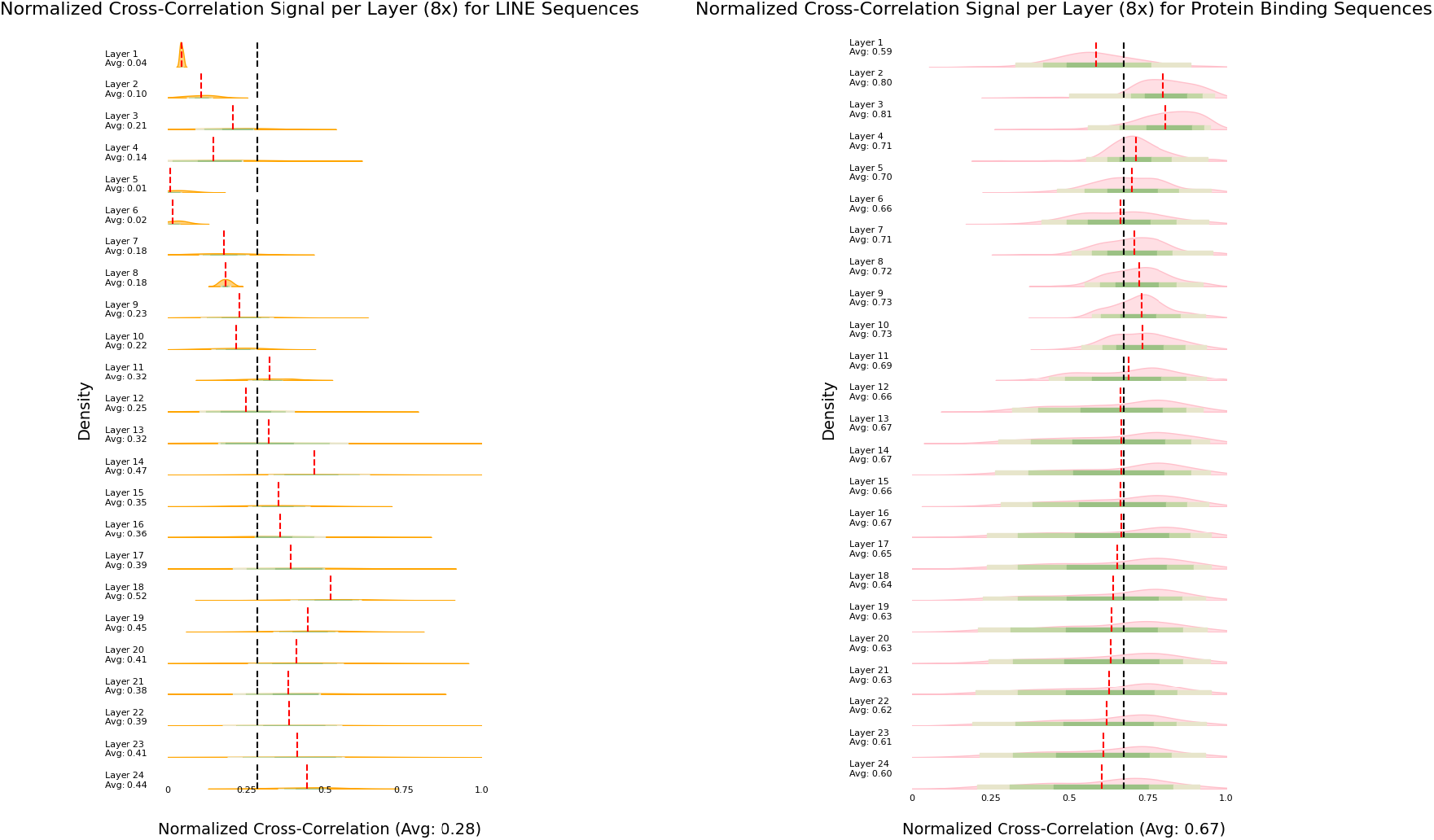
Ridgeline density plot showing distribution of highest correlated features by Normalized Cross-Correlation for subsets of the NCBI RefSeq regulatory element track set from the UCSC genome browser. Comparison between LINE subset (left) and PBS (right). Black dotted lines indicate global average, red dotted lines indicate local layer-wise average. Colored bars represent quartiles.

### Constructing Knowledge Graphs of SAE Activity

Using LLMs to construct knowledge graphs is an increasingly common approach for retrieval-augmented generation (RAG), and presents a potential method for addressing the gap in causal determination for SAE feature meaning during the annotation process [11]. For this task, LLMs are often paired with fine-tuned named entity recognition (NER) or relationship extraction (RE) heads, which are then used to add nodes to a graph and connect them based on their relationships. Applying similar methods to SAE feature activations on a set of sample sequences allows us to explore the activations using graph methods, thereby enabling further analysis of activation patterns and the use of relationship information when comparing sequences to known references.

To construct a SAE knowledge graph as described above, we can apply NTv2 to a sample dataset to generate latents for a given layer of NTv2, parse the feature activations by connecting the sequence IDs to the strongest activating features, and type each edge according to the tokens which caused the activation. Complete information can be found in the supplementary materials, but the basic graph structure is a *typed heterogeneous multigraph* as shown below:

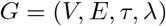

Where:

- *V* is the vertex set
- *E* is the edge set (multiset, allowing parallel edges)
- *τ* : *E* → Σ is the edge type function
- *λ* : *V* ∪ *E* → *A* assigns attributes to vertices and edges

Although the full knowledge graph we constructed is too large to practically visualize, the subgraph in *Figure 3* demonstrates the basic structure of this graph. Using the full collection of RefSeq annotations as found in the UCSC genome browser, we can also record functional metadata for each sequence to perform additional analysis on different sub-communities of SAE activation.

**Figure 3:**
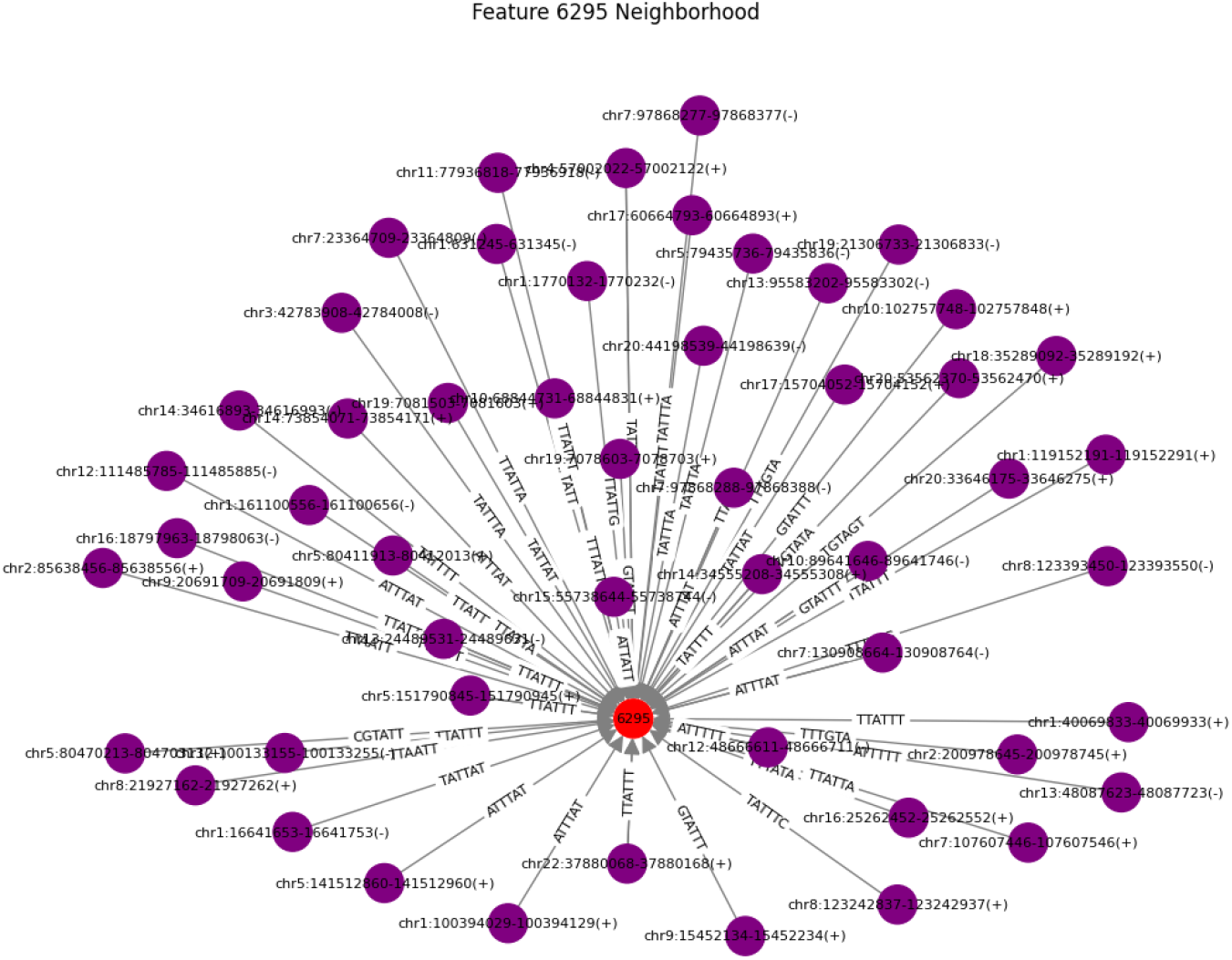
Neighborhood subgraph of feature 7244 containing all inbound edges from first-degree neighbors. The full knowledge graph was constructed from layer 23 SAE activations (10240 feature neurons) from NTv2-500m-human-ref on a set of putative cisplatin binding sequences. Purple nodes represent sequences, edge labels describe the nucleotide sequence of the token, and red nodes represent the central feature of the subgraph.

Our sample dataset comes from work by Krishnaraj and colleagues [12], who are investigating the binding of RNA sequences to the chemotherapy drug, Cisplatin. By developing a novel click-chemistry transcriptome assay called PlatRNA-seq, they identified a subset of transcripts and RNA fragments that have the capacity to bind to Cisplatin, potentially revealing a new mechanism for chemotherapy resistance. These RNAs are believed to form stable rG4 quadruplex structures in the presence of positive cations such as K+, with cisplatin competing in this role prior to initial formation [13]. However, structural characterization in RNA is still somewhat limited, and large RNA structure databases that contain reliably characterized structures of these transcripts of interest are not yet established. Thus, the precise mechanism is still unknown. Using the mapped loci from Krishnaraj and colleagues to obtain DNA reference sequences (GRCh38), we combined this set of sequences with an equivalent set of randomly selected sequences from the remainder of the human reference genome to create training and testing datasets for CNN classification heads as well as knowledge graphs and intervention studies.

The ongoing line of work by Krishnaraj and colleagues provides an interesting opportunity to address a point of discussion from Simon and Zou [7], who suggested that mechanistic interpretability may be able to reveal novel biological patterns learned during pretraining that have not yet been considered by the broader scientific community. This hypothesis suggests that the process of annotating gLM feature activations may reveal novel mechanisms, syntax, or binding motifs which would explain the observations by Krishnaraj and colleagues [12]. At present, methods for addressing this question are extremely limited as most approaches to mechanistic interpretability in genomics require a known reference or ground truth. Graph structures are relational, thereby enabling unsupervised analysis of the graph itself to derive patterns of activation which may indicate functional patterns in the underlying biology.

To demonstrate this, we began by constructing SAE knowledge graphs as described above for the cisplatin-binding sequences and non-cisplatin-binding sequences using the final layer of the NTv2 encoder. Based on earlier observations, features at this layer are likely the most fragmented and granular, thus the likelihood of finding novel patterns is highest in this layer. It is also critical to understand why SAE latents fail to capture currently understood regulatory elements consistently in this layer when the concept was more readily extracted in previous layers.

### Graph Topology

By nature, the sequence-to-feature graph structure is relatively sparse, meaning that most nodes are not connected to each other. This can be represented using a density coefficient that reflects the ratio of existing edges to possible edges, where, a perfectly connected graph has a density of one, and a graph with no connections between nodes has a density of zero. The cisplatin-binding graph had a density of *4*.*01e-3*, and the non-binding graph had a density of *2*.*16e-2*, indicating that most sequences tended to have a few key active features across their length, and most of these were common enough to be useful descriptors of the cisplatin-binding and non-binding phenotypes. Although they are both sparse, the non-binding graph is an order of magnitude more dense than the cisplatin-binding graph, indicating more diverse SAE feature activity the non-binding phenotype.

Ideally, the graph topology should be relatively sparse with key features being major hubs of feature activation. This enables us to perform focused intervention studies and demonstrates the degree of monosemanticity that is being achieved for the target concept. In *Figure 4*, we demonstrate this by progressively removing the highest PageRank centrality feature nodes from the graph and plotting the rate of fragmentation by counting the number of disconnected subgraphs and measuring the density coefficient. PageRank centrality was originally used for web search ranking and measures the importance of nodes by estimating the probability of landing on that node via a combination of random walks and jumps like someone surfing the internet and clicking hyperlinks; these metrics and subsequent graph topological analyses were implemented using the NetworkX library [14]. The cisplatin-binding graph fragments much more readily than the non-binding graph, suggesting that NTv2 is able to represent the cisplatin-binding phenotype with fewer SAE latents than the randomly selected non-binding sequences. The precipitous drop in graph density after removing the first 10 nodes also indicates the highly centralized structure of the graph. The cisplatin-binding graph contains 2071 feature nodes, while the non-binding graph contains 3039. This means that graph density reaches a minimum at approximately 4% and 6.5% of feature nodes removed, respectively.

**Figure 4:**
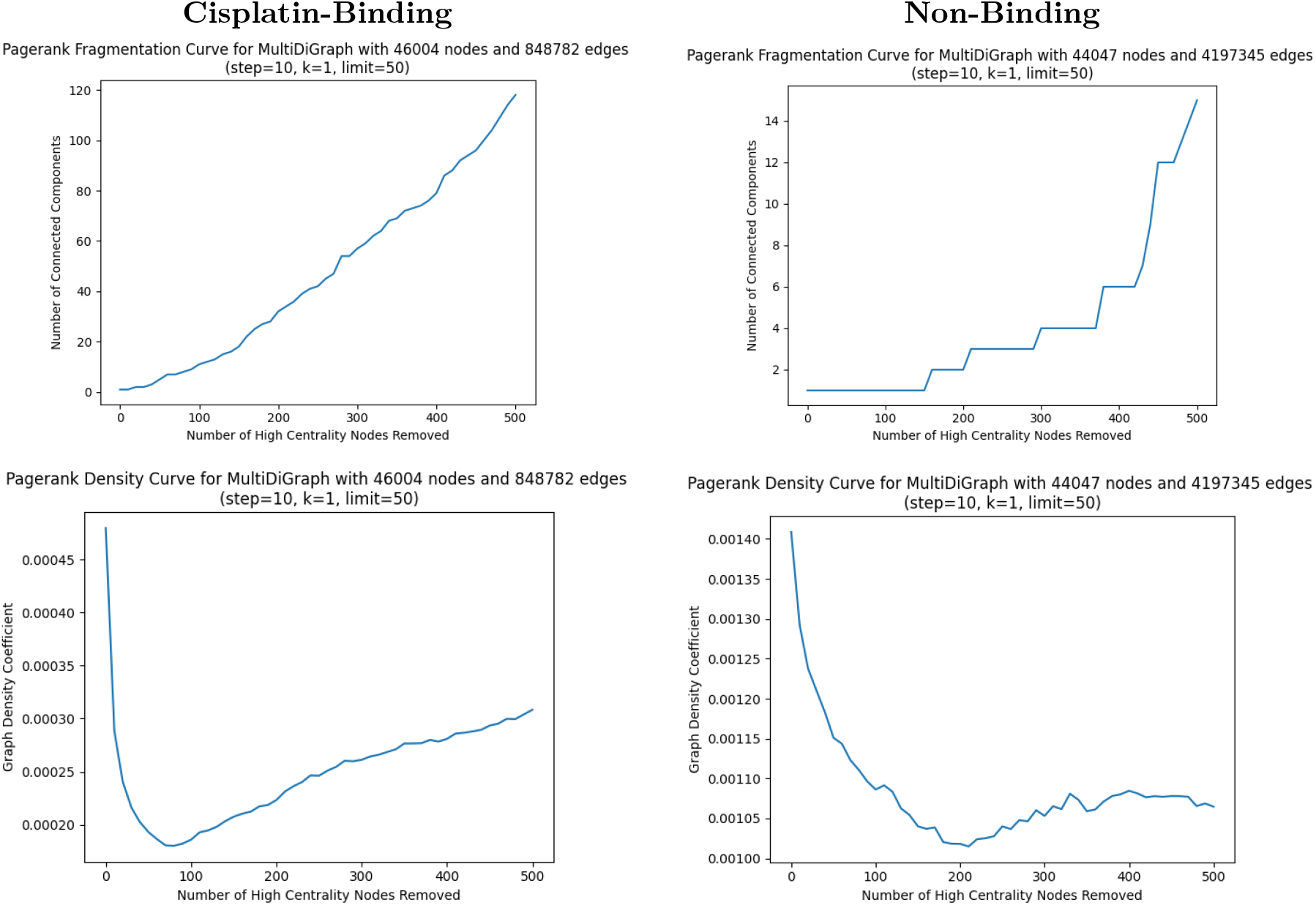
Fragmentation (top) and density shift (bottom) curves for the cisplatin-binding (left) and non-binding (right) graphs when progressively removing the highest PageRank centrality feature nodes. “Number of Connected Components” refers to the number of disconnected subgraphs. These are segregated subgraphs which are connected to themselves, but not connected to other graphs or communities by edges from any of its members.

If we continue to remove nodes from the graphs in this manner until visualization becomes practical, we also find interesting layering patterns as shown in *Figure 5*. With approximately 50% of their feature nodes removed, the cisplatin-binding graph exhibits a layered fragmentation pattern where some sequences are still connected to multiple remaining features, but many of them are fully disconnected and many sequences have been dropped from visualization as complete orphan nodes. Approximately 10.3% of nodes remain in the cisplatin-binding graph, some of which are loosely associated with the core component, while the minor components are spread in a ring-like structure around the expanding core. By contrast, the non-binding graph retains 27.6% of nodes, mostly in the core, and exhibits minimal fragmentation. The retention of a major graph core component after removing the central nodes in this case does suggest some sequences still have very diverse feature sets, and the degree of interdependence observed may be proportional to the capacity of NTv2 to capture the targeted phenomenon.

**Figure 5:**
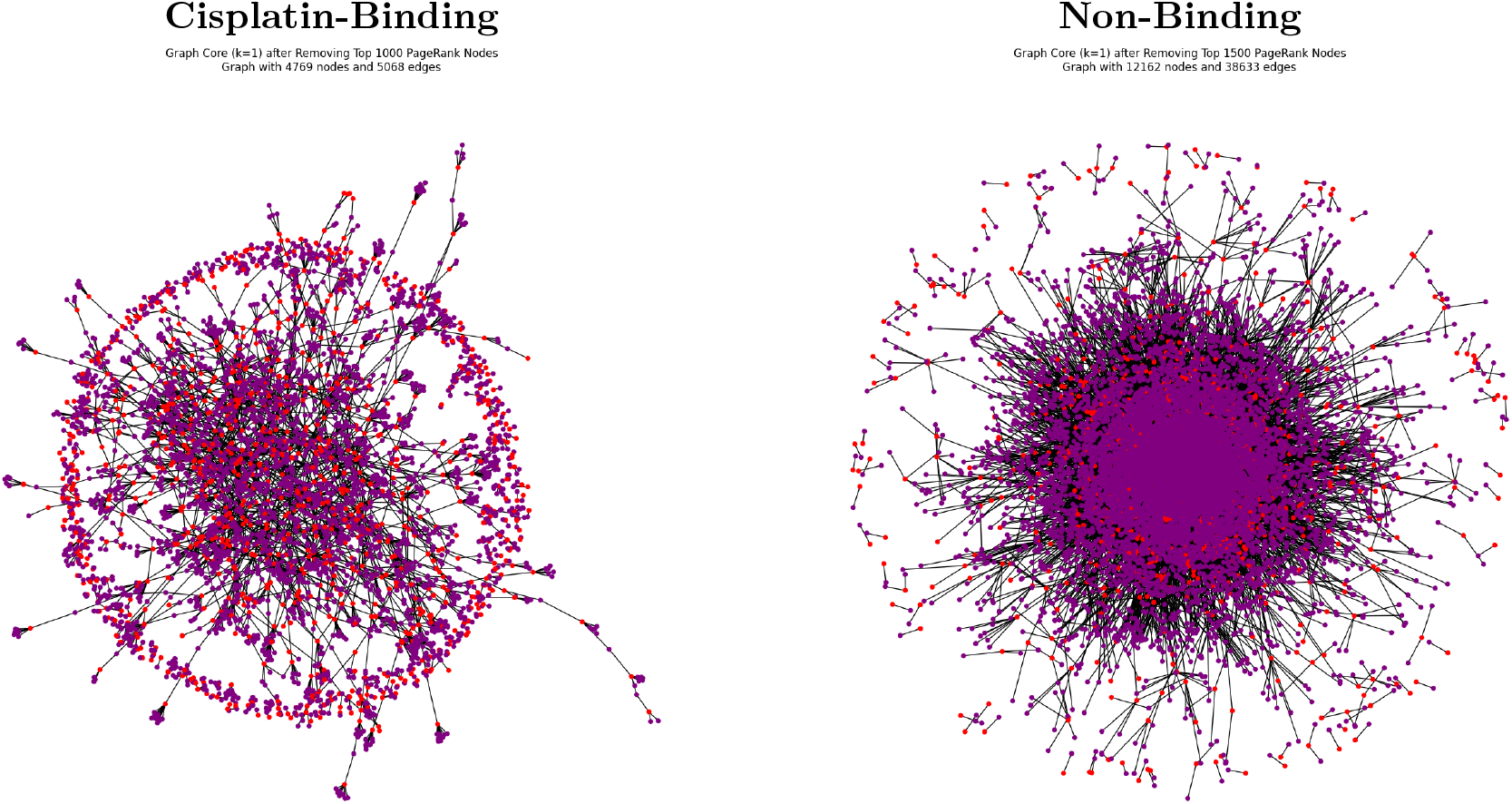
K-core fragmentation visualization of the cisplatin-binding (left) and non-binding (right) graphs, highlighting structural differences between the two phenotypes. Feature nodes are colored red, and sequence nodes are colored purple.

The above observations suggest that the cisplatin-binding sequences have a shared set of key features describing their location within the latent space of NTv2 that we can identify, whereas the non-binding class has diverse feature activation sets that make it suitable for a baseline comparison. Since exposure to cisplatin is usually the result of medical intervention, we do not expect a single latent to represent this non-physiological phenomenon. However, the latents appear to still be able to represent this information using a small set that we can target and search for.

### Selecting Features for Intervention

To identify differentially important SAE features between the two sets of sequences, we compared their centrality in their respective graphs as shown in *Figure 6*. As we did above, we selected PageRank for this purpose since it is an efficient algorithm for large-scale directed graph analysis, and we found it to be the most effective at selecting nodes which would fragment the graph. As with most centrality metrics, a higher centrality score means the feature is more relevant and important to the selected graph and thus its experimental condition.

**Figure 6:**
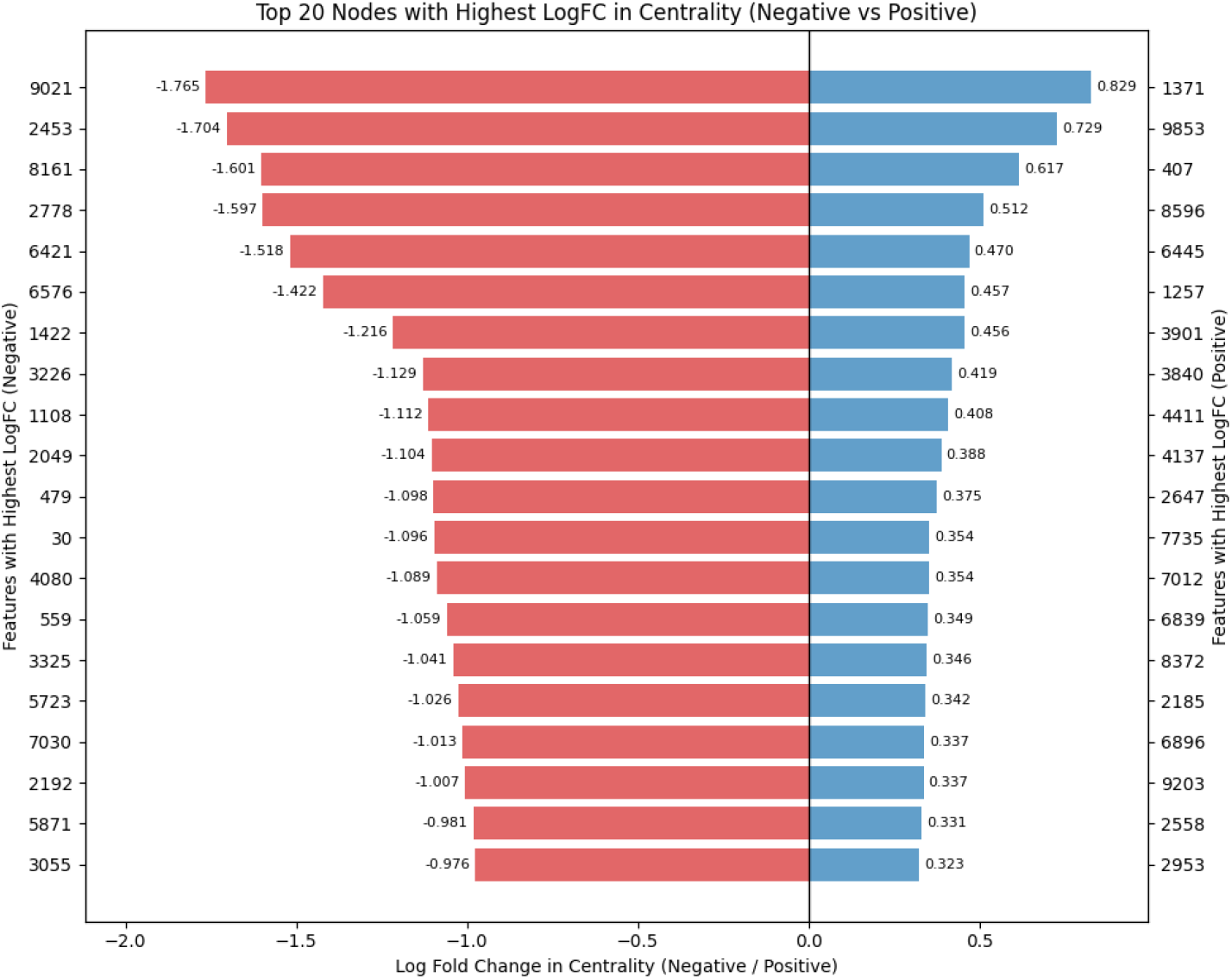
Log-Fold change of PageRank centrality for shared feature nodes between the cisplatin-binding and non-binding graphs. Negative log-fold change indicates the feature had a higher score in the non-binding graph, while positive log-fold change indicates the feature had a higher score in the cisplatin-binding graph. Feature ids are sorted from highest to lowest log-fold change with negatively associated ids on the left y-axis and positively associated ids on the right y-axis.

The differential by log-fold change is important for grounding the feature selection in real effects. While several unique features were identified, their centrality scores were extremely low, suggesting they were more likely to be noise than reliable signal. Using log-fold change also normalizes the magnitude of centrality shifts to avoid biasing toward features that had generally high scores.

### Intervention Studies

The current standard for validating feature annotations is to use the SAE decoder to reconstruct embeddings with a particular feature activated or suppressed. The artificially reconstructed embeddings for the original model can then be used instead of a full forward pass to reliably “steer” gLM behavior. In early work by researchers at Anthropic, subjecting human language models such as Claude to this causes them to generate text reflecting the selected feature, such as sycophancy [6].

Because the generative head of a gLM produces artificial DNA sequences that are themselves un-interpretable by humans, we trained a convolutional neural network (CNN) classification head on NTv2 embeddings to predict whether a sequence would bind or not bind Cisplatin based on the data provided by Krishnaraj and colleagues [12]. The CNN was composed of 2 convolution layers and 2 max pooling layers with a single linear layer out. The convolution kernel was 1-dimensional, sliding across the sequence, while the pooling layers operated on the embedding dimension, gradually reducing the representation down to a logit vector used to compute class probabilities. Using cross-entropy loss, we trained the CNN for up to 100 epochs on shards from an 80:20 train-test data split combining ∼44 thousand binding sequences with ∼40 thousand non-binding sequences. Early stopping was conditioned on decreasing loss over 10 epochs and triggered after 33 epochs, resulting in a model with a test accuracy of 96.2% and an F1 score of 0.9648 after only seeing 33% of the training set. Using the 20% holdout set of test data, we used the unseen sequences to perform steering experiments using the 8x expansion SAE for layer 23 of the NTv2 encoder to create modified embeddings as input to the CNN classification head.

In prior studies, simple clamping of feature values to a given threshold produced desirable shifts in model behavior [6, 7, 8]. However, the therapeutic context of cisplatin exposure is inherently artificial, leading to unexpected probability shifts in the CNN classification head as shown in *Figure 7*. As a result, we found it necessary to experiment with various intervention patterns to elicit the underlying mechanisms. We adopted a grid search intervention protocol involving initial clamping of the selected feature to a minimum value across each token position, followed by element-wise multiplication of an intervention matrix to scale targeted features by a factor *α* and non-targeted features by 1/*α*, as described below. For latent matrix *Z* ∈ ℝ^*n*×*D*^, target feature *f*, minimum activation *δ*, and scale *α*:

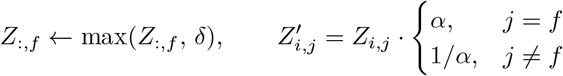

**Figure 7:**
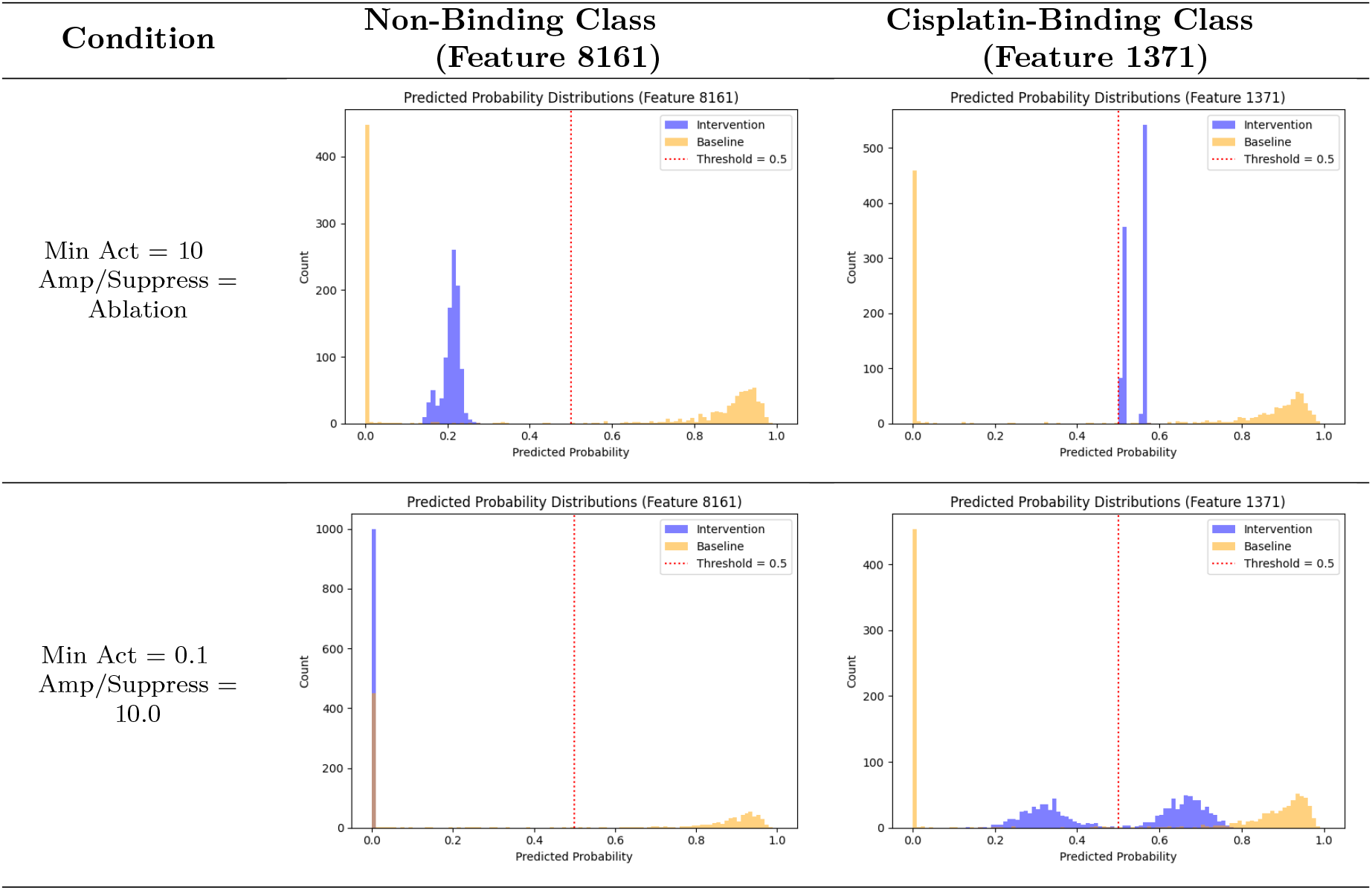
Predicted probability of cisplatin binding for 1000 DNA sequences from the test holdout set, with and without SAE intervention on a selected feature. “Min Act” refers to the minimum activation threshold, and raw feature activation values below this threshold are set to this value for the selected feature. “Amp/Suppress” refers to the target feature amplification factor and the inverse value used to suppress all other feature activity. A value of “ablation” indicates that all non-target features were set to zero. Feature 1371 was associated with cisplatin binding and feature 8161 was associated with the non-binding condition.

A complete formal definition is provided in the supplementary materials. Because of the computational requirements of grid search algorithms, we limited our sequence sample size to the first 1000 sequences from the test holdout set.

#### Predicted Probability Shifts for Selected Features Pre/Post SAE Intervention

As shown in *Figure 7*, the simple clamping of feature activations resulted in consistent shifts in predicted probabilities for all sampled sequences. However, these probabilities often fell between 0.4 and 0.6, well within typical decision-boundary thresholds for binary classification, and not always clearly demonstrating the feature-to-class association identified prior to intervention. One possible explanation is that cisplatin exposure represents a non-physiological perturbation that may be not be directly represented in NTv2 embeddings. Based on this, we altered the intervention to retain some of the original feature information to build a hybrid SAE intervention matrix as described above. Under these new intervention settings, some features produced stronger probability shifts while for other features this same experiment shifted probabilities back toward the baseline.

Supporting evidence for this hypothesis can be found by examining the Area Under the Receiver Operating Characteristic (AUROC) curves for the experiments above as shown in *Figure 8*. The AUROC metric is rank-invariant, meaning that it indicates whether a binding sample has a higher value than a non-binding sample without respect to how large the difference is. As a result, the AUROC curve shifts indicate whether a feature contributes a monotonic shift to all samples, or completely disrupts all predictive signals. Some baseline discriminative signal is present due to the CNN head using zero-padding to account for sequence length in the embedding layer. However, in spite of the noise this introduced, when we use f/1371 to intervene, the discriminative power of the CNN remains, whereas in this experimental setup, discrimination is significantly impeded by intervention on f/8161. This asymmetric AUROC pattern suggests that the non-binding phenotype is represented by distinct and independent inhibitory sequence semantics captured by NTv2, while the binding phenotype is represented by interdependent molecular patterns that must all be present to capture the phenomenon. The asymmetry is consistent with the binding phenotype requiring the co-occurrence of multiple specific sequence features, while the non-binding phenotype is characterized by a single dominant suppressive signal sufficient to override binding predictions — a distinction that mirrors the general principle that disruption of necessary conditions is easier to achieve than synthesis of sufficient ones. This may suggest that the features themselves point to incredibly specific conservation patterns linked to tightly coupled biophysical relationships rather than pathways and functions that may be affected by cell state or environment.

**Figure 8:**
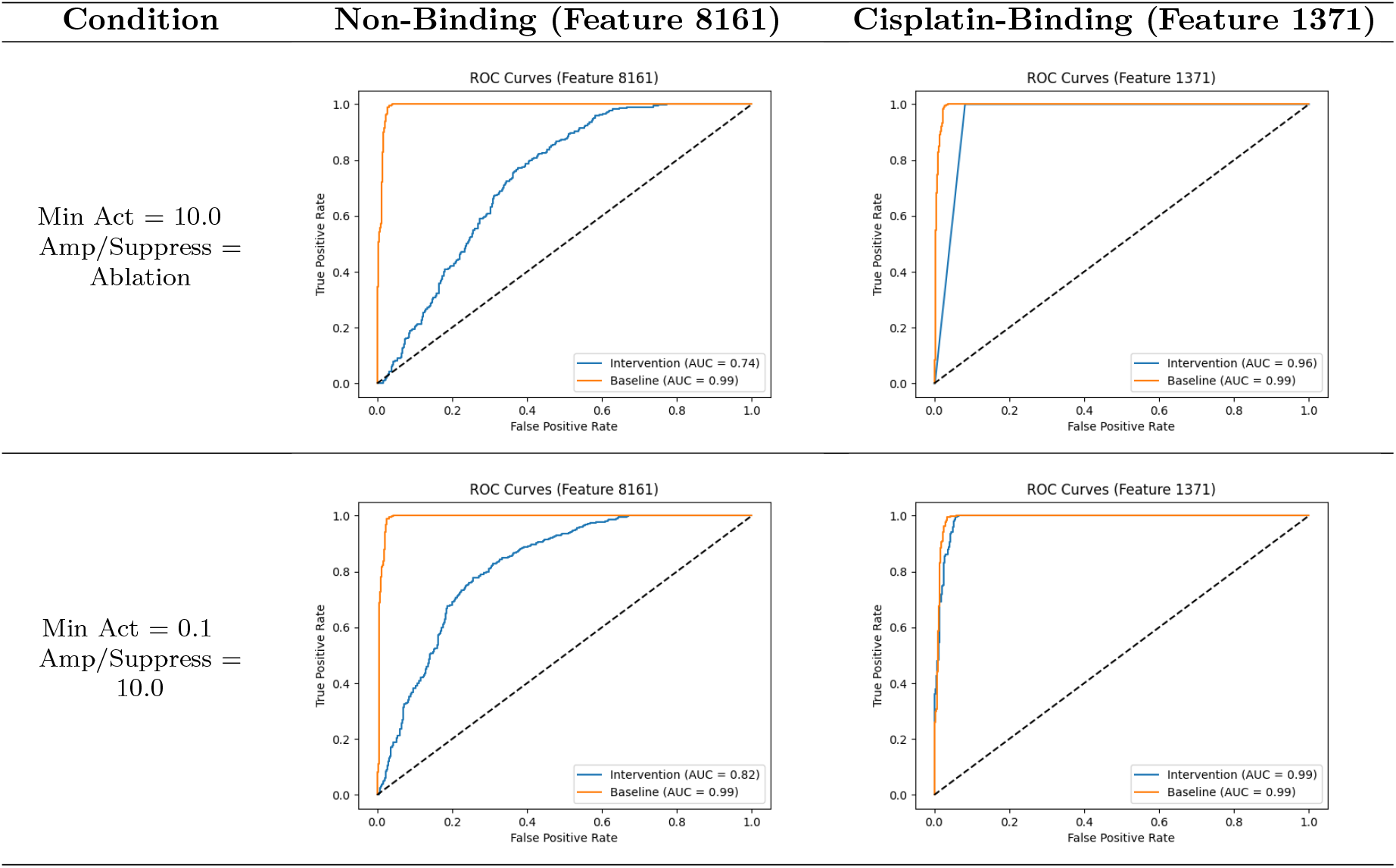
Area Under the Receiver Operating Characteristic (AUROC) curve for 1000 DNA sequences from the test holdout set with and without SAE intervention on the selected feature. Selections mirror Figure 7. “Min Act” refers to the minimum activation threshold, and raw feature activation values below this threshold are set to this value for the selected feature. “Amp/Suppress” refers to the target feature amplification factor and the inverse value used to suppress all other feature activity. A value of “ablation” indicates that all non-target features were set to zero. Feature 1371 was associated with cisplatin binding and feature 8161 was associated with the non-binding condition.

#### AUC-ROC Curves for Selected Features Pre/Post SAE Intervention

### Granularity of SAE Features & Focus on Local Interactions

To gain insight into what these features attend to and how they may be interrelated with the task of predicting cisplatin-binding of RNA transcripts, we adapted dependency mapping techniques from Silva and colleagues [15]. They demonstrated that *in silico* site-directed mutagenesis experiments could identify some functional elements and their dependencies by examining the change in predicted probability for a given base at a given position. We designed a variant of this approach operating on SAE latents, and computed average position-wise deltas of feature activity for all possible substitution variants in selected sequences. The resulting heatmaps in *Figure 9* show how sensitive a feature is at a given base (x-axis) to a substitution at a chosen position (y-axis).

**Figure 9:**
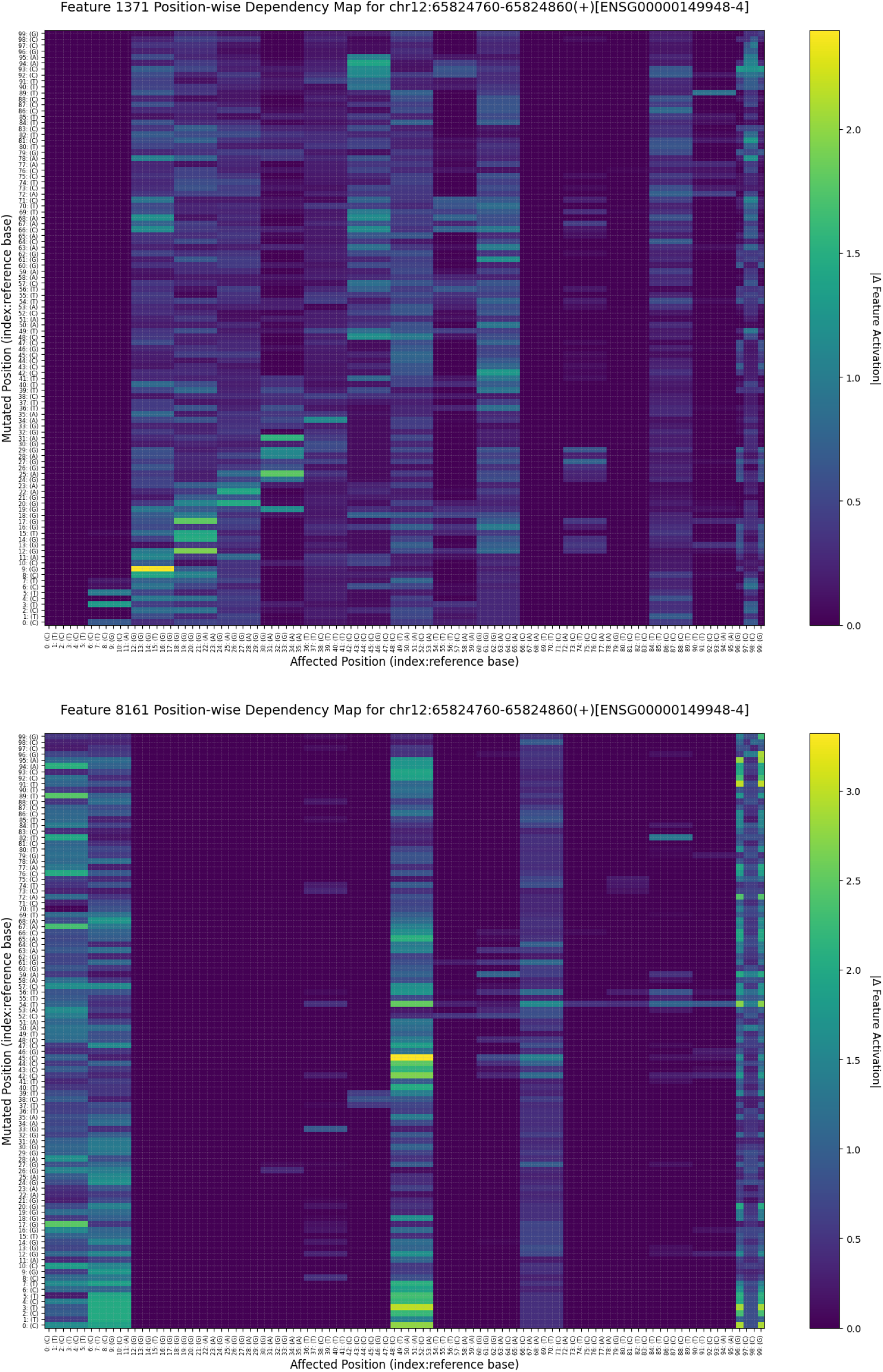
Dependency Maps showing selected feature sensitivity at a given position (x-axis) for substitution at a selected position (y-axis) in a selected cisplatin-binding sequence. Heat color represents the average change in feature activity across all three possible substitutions at the mutated position. Features were selected based on the strongest steering results.

These dependency maps show that even single features can exhibit sensitivity at a surprisingly local level. An interesting point of comparison in *Figure 9* is the relatively local pattern of sensitivity in f/1371 compared to f/8161, which appears to identify key positions that are sensitive to substitutions throughout the sequence. By contrast, changes in f/1371 activity are mostly limited to the token where the mutation occurred. Most of the features we performed intervention on had patterns similar to f/1371, and f/8161 was the only feature to achieve complete leftward probability shift during the hybrid intervention experiment. Combined with recent literature, this suggests that despite the transformer architecture’s advantages in tracking long-range relationships between tokens, the pre-training objective incentivizes gLMs to focus on local biophysical relationships and context over broader regulatory logic that depends on interactions between distant elements.

If these latents accurately approximate the basis of the gLM latent space, it may be nearly impossible for broad regulatory syntax and regulatory pathways to be directly captured by gLM representations that appear to focus on local biophysics. Representing such information would require a composition of several of these latents, at a minimum, or may not be possible to represent at all if the regulatory mechanics are dependent on extrinsic factors such as metabolic feedback loops. If the pre-training objective is the source of the bias, it also remains unclear whether expansion of parameter count or context window size would overcome a bias toward local information.

### Visualizing Feature Sensitivity to Putative Physical Interactions

Although Krishnaraj and colleagues [12] were able to determine that cisplatin binds to putative rG4 quadruplexes and can inhibit cation stabilization by potassium, the exact mechanism is still undetermined. To discuss what that mechanism might be and what biophysical properties these features were sensitive to, we generated predicted structures with Boltz-2 [16] for a series of experimentally validated rG4 quadruplexes in the Quadratlas database [17] that were found wholly within the mapped regions identified by Krishnaraj and colleagues [12]. These structures were then linked to dependency maps of selected features by projecting the dependency maps into a 1-dimensional representation using a spectral graph laplacian to produce a single per-base sensitivity score that could be visualized on the predicted structures generated by Boltz-2.

As we examine these structures, it is important to note that RNA structures tend to be highly dynamic, and some conformations may require interaction with various proteins to become energetically favorable [18]. These predicted structures are valid for hypothesis generation, but are not themselves conclusive. While Boltz-2 has predicted mostly hairpin structures for the selected sequences, many conformations are possible, including varying intermediates, in the absence of experimentally characterized structures.

In the predicted Boltz-2 structures, it is common to observe cisplatin within the major groove of the RNA hairpin double helix as shown in *Figure 10*. This makes intuitive physical sense given it is the most obvious binding site in the predicted structure, and coincidentally aligns with some feature sensitivity scores and known biochemistry [19, 18, 20]. For example, the canonical therapeutic mechanism for cisplatin involves forming intrastrand crosslinks between adjacent purine bases in DNA. Cisplatin is typically aquated when it passes into the cytosol due to the low intracellular chloride concentration, which allows it to form crosslinks via nucleophilic attack, usually by the N7 position on guanine [19]. In *Figure 11*, we see that f/3378 seems to be sensitive to a series of guanine bases which are in exposed positions that would be conducive to this mechanism of cisplatin crosslink formation. Alternatively, zinc finger motifs are commonly found in proteins which bind DNA and RNA for various regulatory functions [18, 20]. These motifs commonly include GC boxes and purine-rich sequences involved in rG4 quadruplex formation, where arginine, histidine, and threonine residues may interact to stabilize and fold the RNA into the quadruplex structure.

**Figure 10:**
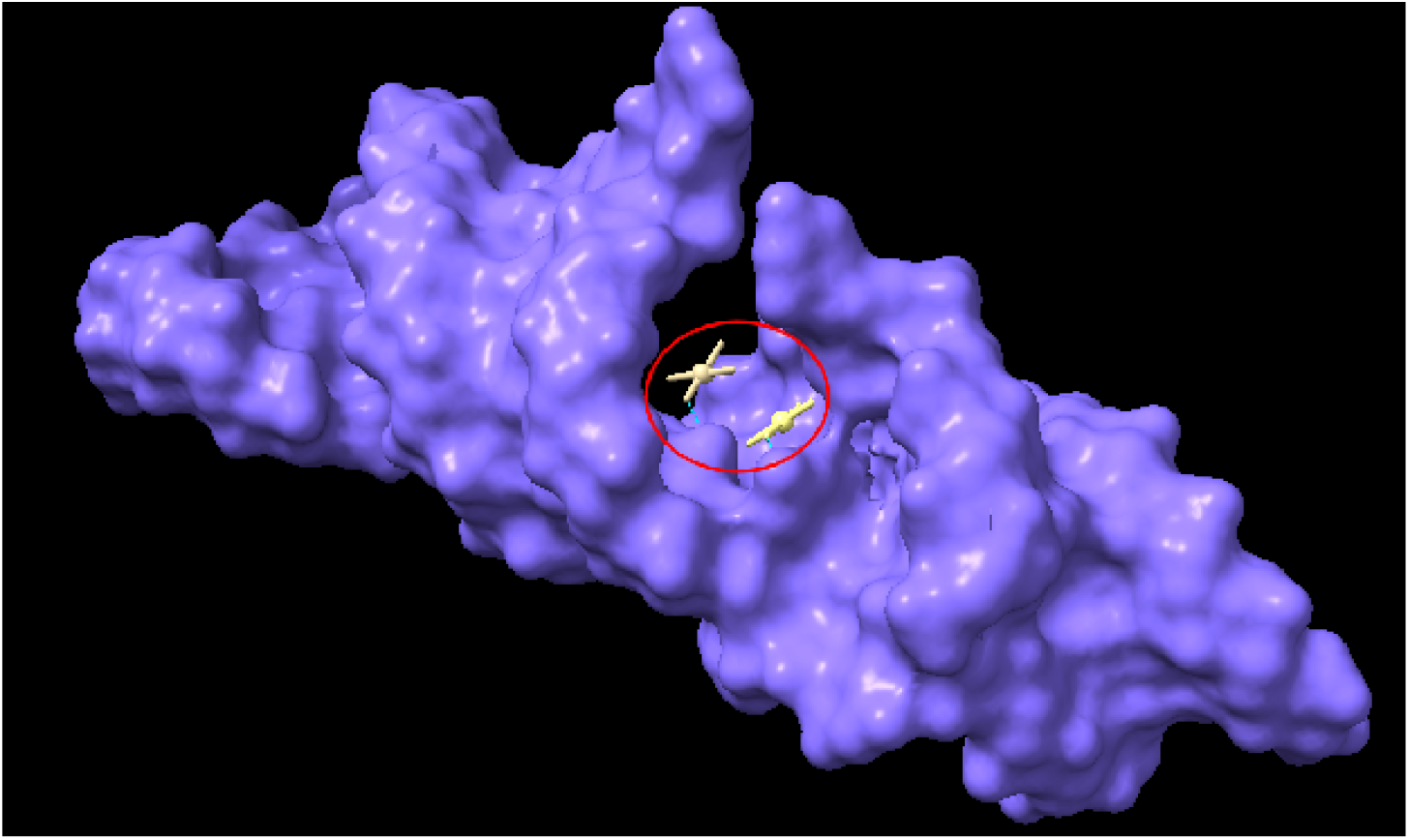
Electron space filling visualization of predicted RNA secondary structure for chr1:31938031-31938149(-) by Boltz-2 in the presence of 2 cisplatin molecules. The red circle highlights the presence of cisplatin (white) within the major groove of the RNA hairpin helix (purple). Dotted blue lines indicate hydrogen bonding.

**Figure 11:**
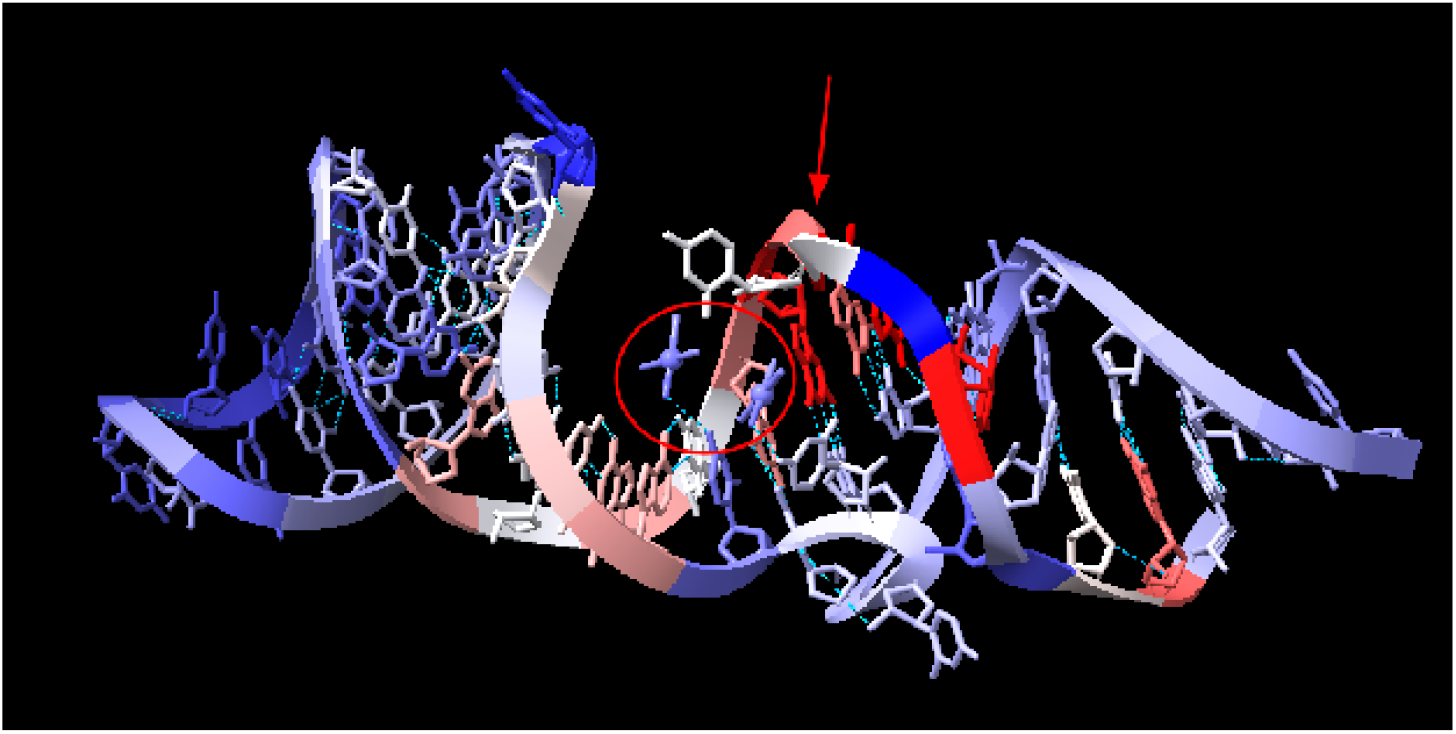
Stick and ribbon visualization of predicted RNA secondary structure for chr1:31938031-31938149(-) by Boltz-2 in the presence of 2 cisplatin molecules. The red circle shows the cisplatin molecules (purple) within the major groove of the RNA hairpin helix. The helix is colored according to feature sensitivity score (f/3378) derived from dependency mapping via NTv2-500m-human-ref and SAE output from the final encoder layer. Red indicates high sensitivity and blue represents low sensitivity. The red arrow points to exposed consecutive guanine bases in the major groove which have notably high f/3378 sensitivity score.

While this evidence should not be construed as empirical evidence for a biochemical mechanism of cisplatin-RNA binding, the existence of plausible biochemical mechanisms which explain f/3378 sensitivity provides a structurally grounded data point that reinforces the pattern emerging from graph topology, intervention asymmetry, and dependency locality — that NTv2 representations encode local biophysical constraints with sufficient fidelity to enable out-of-domain predictions, although targeted experimental validation is still required [21].

## Discussion

Although NTv2 was unable to track many of the current annotated regulatory elements from the reference genome, it appears to have tracked minute biophysical syntax that enabled downstream predictions of non-physiological phenomena. This granular capture of syntax can be understood functionally through the relationship between the pre-training objective, information entropy, and evolutionary constraint. By minimizing cross-entropy loss during pre-training, gLMs are inherently incentivized to compress and predict low-entropy sequence motifs. In the genome, regions of low information entropy correspond to high evolutionary conservation, which often strictly preserves local biophysical interactions and stable secondary structures (e.g., RNA G-quadruplexes).

Consequently, the “minute syntax” encoded by individual SAE features appears to represent these low-entropy, biophysically constrained structural building blocks. To predict a complex event like cisplatin binding of RNAs, downstream predictions must aggregate multiple of these syntactical features rather than relying on a single abstract “binding” latent. This explains why gLMs perform well on tasks grounded in tightly coupled local sequence syntax while struggling to natively respond to large-scale regulatory sequences requiring distributed, context-dependent logic — the precise performance pattern identified empirically by Tang et al. [4], who found that gLMs failed to recognize cell-specific regulatory motifs, and by Boshar et al. [21], who found that gLMs matched protein language models specifically on tasks anchored in conserved molecular properties. The difficulty of encoding distributed regulatory logic is further supported by Seitz and colleagues [9], who demonstrated that capturing regulatory elements dependent on long-range sequence context requires task-specific architectural choices beyond what pre-training alone provides — a challenge our results suggest is rooted in the pre-training objective rather than in model capacity alone.

Our findings contribute a mechanistic understanding of why gLMs perform as they do in each domain, however, they also demonstrate significant limitations for both gLMs and mechanistic interpretation of these models. In terms of the hidden biological patterns among gLM representations, our results are consistent with other studies that have indicated the presence of this information in pre-trained representations [9, 15]. However, current methods are fundamentally limited in determining what the function of these novel elements is, which limits our ability to design studies to validate or investigate the function of these patterns in the genome. There is also a limitation of scale, as current interpretability methods tend to either provide low-resolution insights at scale or high-resolution insights on the scale of individual genes due to the use of algorithms with exponential complexity. Unpacking these hidden biological patterns therefore requires algorithmic innovations that improve on the limitations of current methods.

For mechanistic interpretability in particular, the extraction of SAE latents has already been demonstrated at scale [1], but the annotation and exploration of what those latents mean still has major obstacles to overcome. We demonstrated several innovations that may contribute to addressing those challenges, but they primarily address epistemic limitations and still scale with exponential complexity, meaning they will quickly become impractical for large scale annotation efforts. Further development of interpretability efforts must address the computational complexity challenge to enable widespread adoption.

## Data and Code Availability

All source code is publicly available at: https://github.com/ek775/Hidden-State-Genomics. Large data files and artifacts not pulled from public sources can be found on Zenodo at 10.5281/zenodo.20073353.

## Acknowledgements and Conflicts of Interest

We would like to thank Krishnaraj and colleagues for sharing their PlatRNA-seq data and mapped loci prior to publication and for granting permission to use their data in our work.

## Supplementary Materials

### Supplementary Methods S1: Notation

Let a sequence produce token-level hidden states *h*_*i*_ ∈ ℝ^*d*^ for positions *i* = 1, …, *n*. Let *m* denote the SAE expansion factor and *D* = *md* denote the latent dictionary size. Let *z*_*i*_ ∈ ℝ^*D*^ be the sparse latent activation at position *i*.

### Supplementary Methods S2: Sparse Autoencoder Training Objective

We train a one-hidden-layer sparse autoencoder over transformer hidden states with reconstruction and sparsity regularization. For each token embedding *h*_*i*_, the model produces *z*_*i*_ and reconstruction 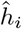. The optimization target is

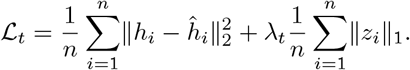

The sparsity weight is linearly annealed during early optimization:

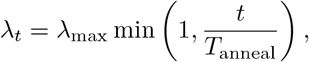

where *t* is the training step and *T*_anneal_ is the annealing horizon. Hidden-state sequences are shuffled, partitioned with a 10% validation holdout, and processed in shards to control memory footprint. Model selection uses validation tracking with early stopping.

### Supplementary Methods S3: Knowledge Graph Construction

We construct a typed heterogeneous directed multigraph

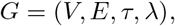

with disjoint vertex sets

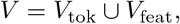

where *V*_tok_ are nucleotide token strings and *V*_feat_ = {0, 1, …, *D* − 1} are feature indices. For each sequence position *i*, define the dominant feature

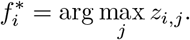

A directed edge 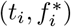 is added from token *t*_*i*_ to feature 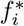. The edge type mapping *τ* assigns token-mediated edge identity, and *λ* stores sequence and genomic metadata (sequence ID, coordinate, strand, and annotation attributes). This yields a multigraph with parallel edges across repeated token-feature events in distinct genomic contexts.

### Supplementary Methods S4: Feature Intervention Operator

For latent matrix *Z* ∈ ℝ^*n*×*D*^, target feature index *f*, minimum activation *δ* ≥ 0, and intervention scale *α* ≥ 0:

1. Threshold clamp on target feature:

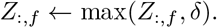
2. Construct multiplicative intervention mask *V* :

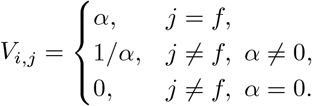
3. Apply intervention:

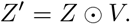

When *α* = 0, non-target features are ablated. Intervened representations are then passed to downstream prediction for comparative evaluation against baseline predictions.

### Supplementary Methods S5: Data Splits and Evaluation Protocol

Cisplatin-binding and non-binding sequence sets are deduplicated prior to partitioning. Data are split into train-test with ratio 80:20, and the training partition is further split into train-validation with ratio 80:20, yielding an effective 64:16:20 train-validation-test composition. Intervention analyses are performed on a reduced holdout subset under compute constraints, as noted in the main text.

### Supplementary Methods S6: Reproducibility Inputs

Core runtime inputs include model reference path, sequence repository path, reference cache files, and cloud storage location for large artifacts. Source data include BED or FASTA sequence inputs, genomic annotations, trained SAE checkpoints, and downstream classifier weights.

### Supplementary Methods S7: CNN Classification Head Architecture and Training

To probe whether SAE representations are predictive in a downstream supervised task, a convolutional neural network (CNN) classification head is trained to discriminate cisplatin-binding RNA sequences from non-binding controls. The classifier accepts as input either the SAE latent matrix *Z* ∈ ℝ^*L*×*D*^ (features mode) or the transformer hidden-state matrix *H* ∈ ℝ^*L*×*d*^ (embeddings mode), where each sequence is zero-padded to a fixed length of *L* = 1000 tokens.

#### Architecture

An adaptive kernel width *k* = max (⌊*D*^1/5^ ⌋,2) is derived from the input feature dimension *D*. The network applies the following operations sequentially:

1. Convolutional block 1: *D* → ⌊*D*/8 ⌋channels, kernel width *k*, half-padding ⌊*k*/2 ⌋, ReLU activation.
2. Convolutional block 2: ⌊*D*/8 → 64⌋ channels, same kernel and padding, ReLU activation.
3. Max-pooling: kernel width *k*, stride 6, ReLU activation.
4. Dropout with rate 0.5.
5. Adaptive max-pooling to temporal dimension 1, ReLU activation.
6. Fully connected layer: 64 → 2 logits for binary classification.

#### Training objective

Parameters are estimated by minimizing categorical cross-entropy:

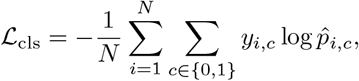

where *y*_*i,c*_ ∈ {0, 1} is the one-hot class indicator and 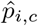 is the softmax-normalized predicted class probability for sample *i* and class *c*.

#### Optimization

The Adam optimizer is used with learning rate *η* = 10^−3^. Training data are uniformly fragmented across epochs, allowing the full dataset to be traversed over the training run without loading all upstream representations into memory simultaneously. Early stopping against validation loss terminates training when no improvement is observed within a fixed patience window.

